# Multifaceted impact of specialized neuropeptide-intensive neurons on the selective vulnerability in Alzheimer’s disease

**DOI:** 10.1101/2023.11.13.566905

**Authors:** Manci Li, Nichole Flack, Peter A. Larsen

## Abstract

**INTRODUCTION:** Widespread disruption of neuropeptide (NP) networks in Alzheimer’s disease (AD) and disproportionate absence of neurons expressing **<u>h</u>**igh **<u>N</u>**P-**<u>p</u>**roducing, coined as HNP neurons, have been reported for the entorhinal cortex (EC) of AD brains. Hypothesizing that functional features of HNP neurons are involved in the early pathogenesis of AD, we aim to understand the molecular mechanisms underlying these observations.

**METHODS:** Multiscale and spatiotemporal transcriptomic analysis was used to investigate AD-afflicted and healthy brains. Our focus encompassed NP expression dynamics in AD, **<u>AD</u>**-associated **<u>NP</u>**s (ADNPs) trajectories with aging, and the neuroanatomical distribution of HNP neuron.

**RESULTS:** Findings include that 1) HNP neurons exhibited heightened metabolic needs and an upregulation of gene expressions linked to protein misfolding; 2) dysfunctions of ADNP production occurred in aging and mild cognitive decline; 3) HNP neurons co-expressing ADNPs were preferentially distributed in brain regions susceptible to AD.

**DISCUSSION:** We identified potential mechanisms that contribute to the selective vulnerability of HNP neurons to AD. Our results indicate that the functions of HNP neurons predispose them to oxidative stress and protein misfolding, potentially serving as inception sites for misfolded proteins in AD.

## Introduction

Alzheimer’s disease (AD) is a neurodegenerative disorder characterized by progressive cognitive decline and memory loss.^1^ The neuropathological features of AD encompass both “positive”—Aβ plaques and tau tangles, glial responses, and cerebral amyloid angiopathy—and “negative” lesions, such as the loss of neurons and synapses.^2^ The majority of AD cases occur sporadically, with no clear understanding of their cause or pathogenesis.^1^ “Epicenters” are described as the site exhibiting the peak pathological changes or atrophy within the brain, often considered coinciding with the initial site of disease onset by network-based degeneration/spread hypothesis for AD.^3,4^ Despite extensive research, gaps in our understanding remain, particularly concerning the selective vulnerability of specific cell types and brain regions to protein misfolding.^5,6^ Investigating these enigmas is crucial for developing targeted interventions and advancing our knowledge of AD’s pathophysiology.^7^

We recently showed the widespread disruption of neuropeptide (NP) networks in AD and the disproportionate absence of cells, mostly neurons, expressing higher levels and diversities of NPs (high NP-producing/HNP cells/neurons) in the entorhinal cortex (EC) of AD brains.^8^ As steps involved in NP production are energy-intensive and HNP neurons dedicated more transcription power to NP production, we hypothesized that HNP neurons face greater energetic demands and higher metabolic stress during aging.^8^ However, the functional characteristics of HNP neurons and the relationship between NP co-expression and other established pathological features of AD are unclear. Although the expression of AD-associated NPs (ADNPs) decreased in the hippocampus during natural aging,^8^ the role of HNP neurons co-expressing ADNPs in influencing cognitive functions and mediating regional brain vulnerability is yet to be elucidated.

In this context, we hypothesize that HNP neurons co-expressing ADNPs may represent cellular substrates implicated in the “epicenters” of cascading protein misfolding events that are the hallmarks of AD (Figure 1).^4–6^ By analyzing single-cell RNA-seq data from the entorhinal cortex and spatiotemporal RNA-seq data, encompassing 1890 human brain samples, we found that HNP neurons had increased metabolic demands as well as increased expression of genes associated with tau misfolding, which was additionally indicated by neurons expressing NPs in AD showing distinct molecular signatures of protein misfolding. We further report that the decrease of ADNP expression with aging was specific to early AD-impacted brain regions and that HNP neurons co-expressing ADNPs were heavily distributed in “epicenters”. These results support the hypothesis that the functional demands of HNP neurons likely predispose them to oxidative stress during aging, perhaps contributing to their early demise in AD.

**Figure 1.**
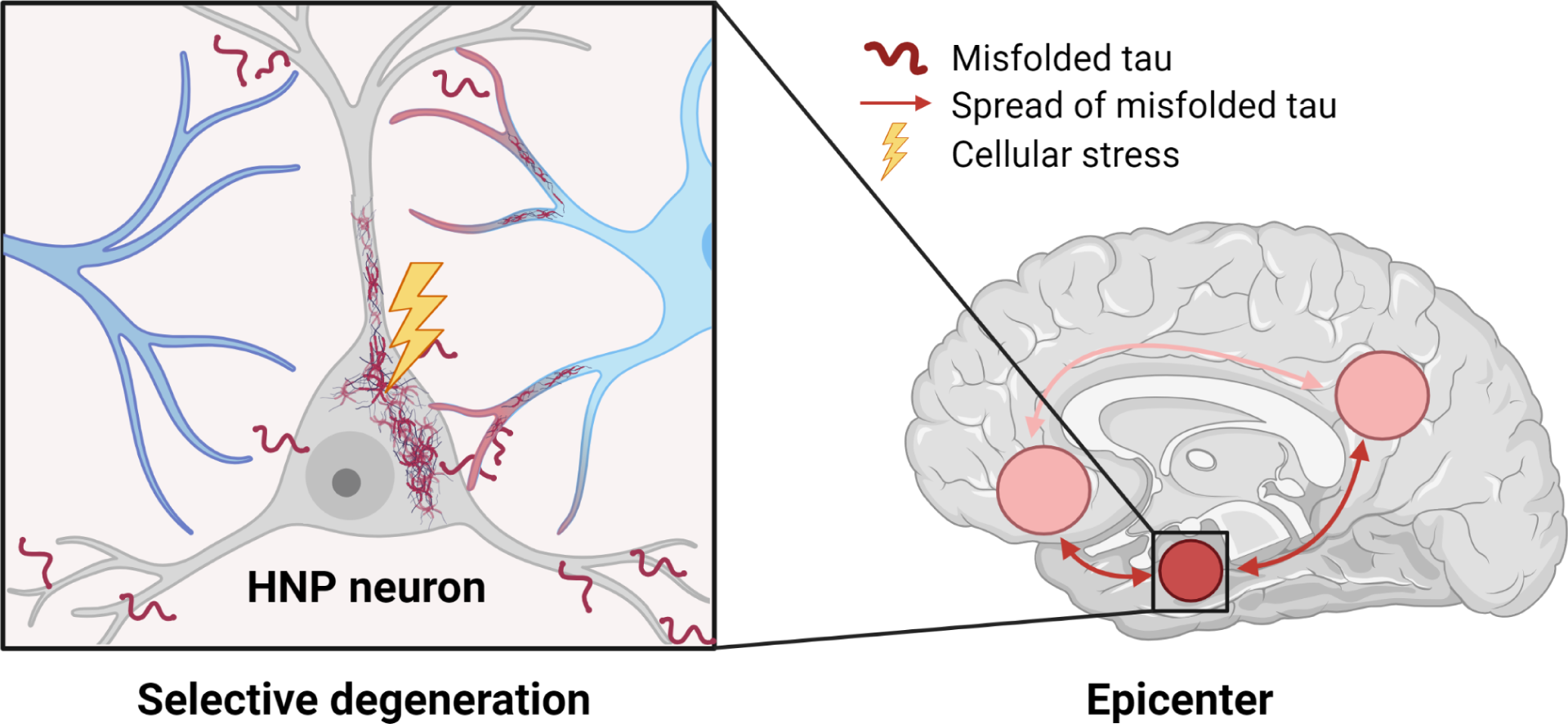
Hypothetical model illustrating the potential mechanisms underlying the selective vulnerability of high neuropeptide-producing (HNP) cells and their contribution to region-specific emergence of Alzheimer’s disease (AD). The model proposes that the unique functions of HNP neurons make them more prone to stress and protein misfolding, and this susceptibility becomes more evident with advancing age. The regional vulnerability observed in Alzheimer’s disease can be attributed to the distribution and density of these cells, as well as their excretory functions and interactions with other brain networks. Specifically, it predicts that (1) temporal, limbic, and prefrontal cortical regions have a higher density of HNP neurons expressing ADNPs; (2) disruption of cellular processes in HNP neurons leads to a decrease in ADNPs during aging and localized formation of misfolded proteins in AD, causing various degrees of cognitive decline and selective degeneration of these neurons; (3) the dynamic paracrine and secretory activities of HNP cells facilitate the propagation of misfolded proteins and transneuronal degeneration in adjacent temporal, limbic, and prefrontal cortical regions, resulting in widespread deposition of misfolded tau in AD.

## Methods

### Acquisition and processing of transcriptome data

#### Single-cell RNA-seq data from the human entorhinal cortex (EC)

Two sets of single-cell EC transcriptome data were included for analysis in this manuscript.^9,10^ Data generated by Grubman et al from six control (CT) and six AD donors (12 total) were obtained.^9^ The detailed documentation of data acquisition, preprocessing steps, and downstream analyses (dimension reduction and cell identification) can be found in Li and Larsen (2023).^8^ Briefly, cells were filtered based on gene expression and mitochondrial content, retaining those with 200-2500 expressed genes and less than 5% mitochondrial reads. Data normalization followed Seurat’s (version 4.1.1; R 4.2.3 unless specified) guidelines using a count per million (CPM) matrix.^11^ Using cell identification methods from *BRETIGEA* and Grubman et al, six primary cell types were identified: astrocytes, microglia, neurons, oligodendrocytes, oligodendrocyte precursor cells, and endothelial cells. Cells were labeled by their highest association score, but some were categorized as unidentified or hybrid based on specific criteria described in detail by Grubman et al as well as Li and Larsen.^8,9^ Only cells identified as neurons were used in the downstream analyses.

In addition, preprocessed and annotated single-cell transcriptome data from human EC generated by the the MIT ROSMAP Single-Nucleus Multiomics Study (MIT_ROSMAP_Multiomics) was downloaded from ADKnowledge portal.^12^ Three diagnostic categories from the ROSMAP study were included in the presented study: control (CT or NCI: no cognitive impairment), mild cognitive impairment (MCI: no other condition contributing to CI), and AD (Alzheimer’s dementia: no other condition contributing to CI (NINCDS/ADRDA Probable AD)).^12^ The clinical study design and detailed diagnostic criteria were described in the original ROSMAP manuscript and deposited on ADKnowledge portal.^12,13^ Individuals with inconsistent clinical diagnosis, clinical cognitive diagnosis summary, and final consensus cognitive diagnosis documented in the ROSMAP study were excluded from the analysis. Methods for isolation of nuclei from frozen post-mortem brain tissue, droplet-based snRNA-seq, and snRNA-seq data preprocessing are available in detail on ADKnowledge portal.^12^ De-identified metadata for individuals and experiments included in this study were detailed in supplementary materials (Table S1). To provide equal representation of each condition, we randomly selected eight AD and CT samples (set.seed = 123) to match the eight available MCI samples. Overall, 41373 neurons (n_neuron = 41373) from eight AD (n_neuron = 16214), MCI (n_neuron = 11732), and CT (n_neuron = 13427) EC regions were included in the final analysis.

#### Spatiotemporal bulk RNA-seq data from human brains

RNA-seq transcript matrices (transcript per million (TPM)) of 11 human brain regions from a cohort of individuals from the general population were obtained from the GTEx portal (Table S2).^14^ Samples from individuals that lacked complete metadata regarding age, sex, or death classification were excluded, as were those that scored 3 or 4 on the Hardy Scale that indicates intermediate or slow death. Only samples with an RNA integrity number (RIN) larger than 6 were included in the analysis. The results were visualized with *cerebroViz* (version 1.0; R 3.6.3)^15^ and BioRender.

### Downstream analyses for single-cell transcriptome data from human EC

#### Correlational analysis of NP transcripts and ADNPs co-expression

The Spearman rank correlation was utilized to assess the relationship between the levels of transcripts of NP and the number of co-expressed ADNPs using *cor.test* in R for both human single-cell transcriptome data. This test is suitable for datasets that do not distribute with normality. Correlation coefficient (rho) and p-values were reported. The significance cut-off was set at 0.05.

#### Differential gene expression (DGE)

DGE analysis was performed for the single-cell dataset generated by Grubman et al. (2019) ^9^ The number of co-expressed NPs was used as a proxy for transcript levels of NPs; neurons were divided into low (0-1), medium (2-5), and high (6+) NP-producing groups (LNP, MNP, HNP) based on the number of co-expressed NPs in both conditions. DGE analysis was implemented between LNP and HNP in control neurons as well as MNP groups in control and AD, using *FindMarkers* function in *Seurat*.^11^ Default parameters for DGE in *Seurat* were used (Wilcoxon Rank Sum test); statistical cut-off was set at 0.05 for a false discovery rate (FDR) adjusted by Benjamin-Hochberg (BH) method.

#### Functional enrichment analysis

The output of the DGE analysis from *Seurat* was used as input to the STRING database (version 11.5).^16^ Key enrichment output from STRING analysis was visualized using the *Enrichplot* package (version 1.18.4).^17^ STRING uses Bonferroni method to correct for multiple comparisons and provides adjusted p-values.^16^ The significance cut-off was set at 0.05 for FDR.

#### Regression analysis of NP transcripts and ADNPs co-expression

Regression analysis was used to discern the relationship between gene transcript levels and the presence of NP. Utilizing the *glm* function in R, general linear models were constructed for each differentially expressed genes (DEG) with transcript levels as the response variable and the number of co-expressed NPs as the explanatory variable.The BH method was used to adjust for multiple comparisons for all p-values. 0.05 was used as the significance cut-off for adjusted p-values. The goal was to identify genes whose expression is notably influenced by the increased number of co-expressed NPs. Results can be found in Table S3.

#### Hypergeometric test of cell abundance and ADNP co-expression

The hypergeometric test was employed using the *phyper* function in R to assess the overlap of genes with decreased expression in AD compared to those that are functionally enriched in HNP neurons.^18^ Specifically, it was defined: 1) the number of overlapped genes as “successes” in our sample (x = 25); 2) genes with significantly decreased expression in AD MNP neurons as “successes” in the population (m = 91); 3) the total number of unique genes expressed by AD MNP and control HNP neurons, minus those with decreased expression in AD MNP neurons, as “failures” in the population (n = 19430); 4) genes with significantly increased expression in HNP neurons as the sample size (k = 307). Since phyper is a cumulative distribution function, the conduction of a one-tailed such analysis (lower.tail = FALSE) would calculate a probability of observing as extreme and more extreme results in the direction of higher values (p-value).^18^ The significance cut-off was set at 0.05.

#### Stratified assessment of cognitive condition-neuropeptide abundance and independence

Using the MIT_ROSMAP_Multiomics dataset, Jonckheere-Terpstra test from *DescTools* package^19^ was used to check if there was a statistically significant trend across the continuum from CT, through MCI, then to AD for both top 50% and top 25% of all ADNP-coexpressing neurons. 0.05 was used as the significance cut-off. Robust t-test (Yuen Two Sample t-test) with a trim of 0.1 in R^19^ was used as a post-hoc test to compare between conditions; 0.1 was used as the significance cut-off to ensure all relevant findings were captured. For each condition in the MIT_ROSMAP_Multiomics dataset (CT, AD, or MCI),^10,12^ The number of co-expressed key NPs (output from ML models as reported previously^8^) was also recorded, ranging from 0 to 3. To investigate the relationships between the conditions and the number of co-expressed NPs, a series of Chi-squared tests to assess the independence between each pair of conditions (AD vs CT, AD vs MCI, and CT vs MCI) were conducted in R. These tests were stratified by the number of co-expressed NPs to examine how this variable might influence the relationships between the conditions. A significance level of α = 0.05 was used for all tests.

### Spatiotemporal correlation analysis between NP gene expression and age

Gene lists of NPs and ADNPs were downloaded from existing publications.^8,20^ TPM count matrix of spatiotemporal RNA-seq data and metadata of human brains were downloaded from GTEx.^14^ All TPM counts were log2 transformed. The correlation between ADNP expression and age was determined using *PResiduals* package,^21^ adjusting for RIN and Hardy Scale. The significance cut-off was set at 0.05.

### Single-cell examination of ADNP-coexpressing HNP (AHNP) neurons across microdissected brain regions

A comprehensive analysis of AHNP occurrences and cell counts across microdissected brain regions was performed using single-cell data and an annotated cell cluster (cluster_annotation) file generated by Siletti et al.^22^ File documentation from the study was downloaded from github link provided by Siletti et al.^22^ Firstly, the presence of ADNPs from the file was quantified and counts for NPs were generated further to calculate non-ADNPs. Based on these counts, we defined ADNP-HNP (AHNP) neurons as those tagged with > 5 ADNPs and < 3 non-ANDPs, accounting for the difference in the input NPs.^8,22^ Concurrently, we estimated the number of cells in different brain regions and dissections based on percentage data extracted from the cluster_annotation file. These estimates were summed across unique regions and dissections to provide a granular view of cell distribution. Additional analyses were conducted on specific regions for MEC, where the dataset was grouped by various attributes such as Neurotransmitter, Subtype, and MTG label,^23^, and the number of cells in each group was summed and visualized.

## Results

### Alterations of HNP neuronal abundance and functions in AD: overlap of HNP dysfunction and molecular signature of AD

Analyzing the single-cell dataset by Grubman et al,^8,9,24^ we showed that the correlation between transcript abundance and the number of co-expressed NPs generally exists for neurons in both control and AD (Figure 2A). Applying the number of co-expressed NPs as a proxy for transcript levels of NPs, we stratified neurons into different groups in AD and controls based on the number of co-expressed NPs, including low (0-1), medium (2-5), and high (6+) NP groups. We observed a similar absence of HNP neurons in AD (Figure 2B; Table S4).^8^

**Figure 2.**
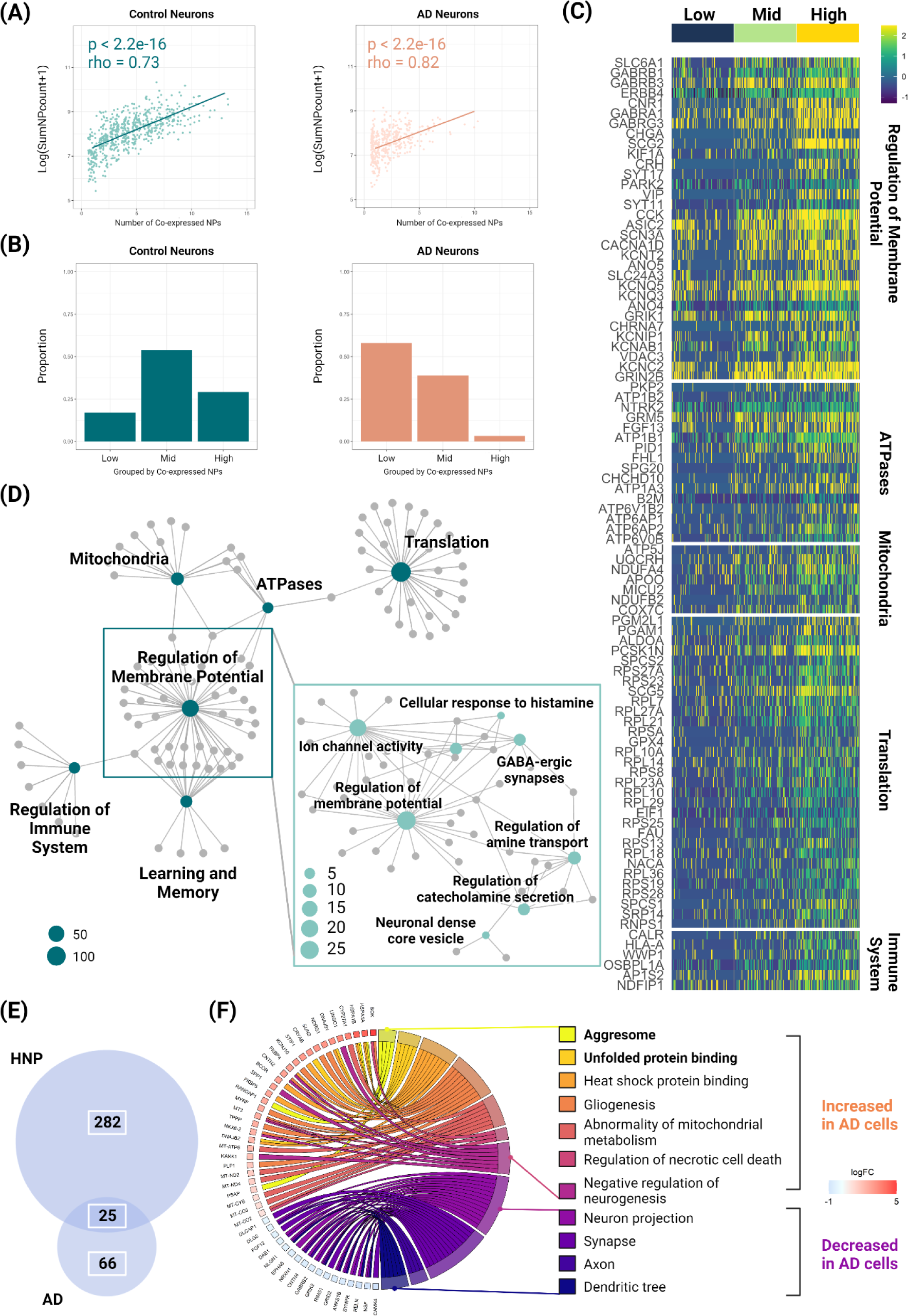
Changes in high neuropeptide-producing (HNP) neuronal abundance and function: overlap of dysfunction and AD molecular signatures. **(A)** The relationship between transcript abundance and the number of co-expressed neuropeptides (NPs) in neurons from both control and AD EC. **(B)** The distribution of neurons based on the number of co-expressed NPs. low: 0-1; medium (mid): 2-5, and high: 6+. Proportion = in-group neuron count in the condition/total neuron count in the condition. **(C)** Heatplot showing differentially expressed genes (increased) in HNP neurons (neurons in high NP co-expression group). **(D)** Gene network plot showing results of functional enrichment for HNP neurons. **(E)** Venn diagram showing the overlap between gene expressions higher in HNP neurons (neurons in high NP co-expression group that express 6+ NPs) and significantly decreased in AD neurons co-expressing 2-5 NPs (MNP neurons). The hypergeometric test was applied to evaluate the overrepresentation of genes upregulated in control HNP neurons but notably reduced in AD MNP neurons; α = 0.05. **(F)** Molecular signatures differentially increased and decreased in AD MNP neurons.

We previously reported a significant reduction of HNP neurons expressing more neuropeptides in the EC of AD brains. As we hypothesized that HNP neurons would experience greater energetic demands and higher metabolic stress, we predicted that HNP neurons would show more metabolic activity. To test this, differentially expressed genes (DEG) were examined between high and low groups of control neurons followed by enrichment analysis.^8,16^ We found that all the NPs expressed at significantly higher levels in HNP neurons were AD-associated NPs (ADNPs),^8^ and genes required for NP transportation, translation, and metabolic processes were significantly increased in HNP neurons in comparison to LNP neurons (Figure 2C&D; Table S5&6). Although it is widely known that GABA is often co-expressed with NPs,^25^ we observed that HNP neurons also participated in other chemical communications, such as histamine and catecholamines, to regulate membrane potential (Figure 2D; Table S5). In addition, ∼36% of DEGs regulating membrane potentials were functionally enriched for learning and memory (Figure 2D; Table S5). Surprisingly, the regulation of innate immune response was increased in HNP cells (Figure 2C&D; Table S5).

To elucidate the increase observed across the LNP, MNP, and HNP groups, we sought to identify genes whose expression was significantly influenced by the level of NP co-expression, which also correlated with the abundance of NP transcripts. We utilized regression models to examine the relationship between gene expression and NP co-expression then ranked the increased DEGs in HNP neurons by coefficients and R^2^. Excluding NP components, genes ranked in the top 10 for coefficients or R^2^ included several long noncoding RNAs, components of A-to-I RNA editing, and *NPAS3*–encoding for a transcription factor related to circadian rhythm regulation^26^ (Table S3; Figure S1). Notably, *ERBB4*, the protein products of which respectively induced tau hyperphosphorylation^27^ was among the top genes related to NP co-expression (Table S3; Figure S1).

Since we discovered that HNP neurons exhibited higher performance in several expected cellular functions–including transportation, translation, and metabolic processes–and participated more heavily in other chemical communications, regulation of innate immune response, and circadian rhythm, we wondered whether dysregulation of processes in these functions may lead to the loss of neuronal functions and the accumulation of tau pathology, which are hallmarks of several neurodegenerative disorders including AD.

Speculating that disrupted functions of neurons expressing more NPs are associated with protein misfolding, we examined the DEGs for neurons stratified by the number of co-expressed NPs and analyzed the neurons in the medium group (as HNP neurons were virtually absent in AD brains). Enrichment analysis revealed that these cells displayed molecular characteristics related to both “positive” and “negative” neuropathology in AD.^2^ We note the decreased DEG in AD cells were significantly enriched for those functionally increased in HNP cells (p < 0.00001; Figure 2E; Table S6&7), indicating that loss of HNP functions participates in the molecular pathogenesis of AD. Genes with protein products showing significantly decreased expression included those with functional roles in axons, synapses, and dendrites (Figure 2F; Table S7). Increased molecular processes included those known to be disturbed in AD, such as negative regulation of neurogenesis, gliogenesis, and abnormal mitochondrial metabolism (Figure 2F; Table S8). Notably, genes involved in forming aggresomes and unfolded protein binding were highlighted,^28^ indicating the active occurrence of protein misfolding in AD cells that co-express NPs (Figure 2F; Table S8).

### Loss of ADNP expression participates in early pathogenesis of AD

Aging is the predominant risk factor for developing AD^1^. The decline in cognitive function during natural aging bears resemblance to that seen in the early stages of AD.^29,30^ Mild cognitive impairment (MCI) is defined as a stage between normal age-related cognitive changes and pathological cognitive impairments, and it frequently precedes and is considered a potential early sign of Alzheimer’s disease.^13^ Therefore, examining datasets from both natural aging and MCI would provide a compelling framework to assess whether disruptions in ADNP expression play a role in the early pathogenesis of AD. Our recent report indicated that ADNP expression decreased with age in the hippocampus;^8^ however, we do not yet know if the decline of NPs with aging is ADNP-and brain region-specific. To strengthen the evidence supporting the early involvement of ADNPs in AD development, we 1) analyzed bulk RNA-seq data from all brain regions with sufficient sample size generated by GTEx and 2) assessed the abundance of HNP neuron co-expressing ADNPs in the EC of AD, MCI, and control (CT) brains using recent single-cell transcriptome data.^10^

With respect to aging, if our hypothesis were to hold, only brain regions affected by early AD should show age-related changes for ADNP expression and only the accumulative expression of ADNPs, but not other NPs, should decrease with age in the human brain. We first examined the expression of ADNPs during aging among 11 brain regions selected from GTEx (Table 1).^8,14^ Supporting our hypothesis, we found that only the hippocampus, frontal cortex, anterior cingulate gyrus, and amygdala—all of which are brain regions affected by early AD^31–36^—showed a significant decrease of ADNP transcription during aging among 11 brain regions selected from GTEx (Table 1; Figure 3A).^8,14,37^ We also comprehensively examined the expression of NPs in these brain regions and whether the expression of an individual NP was correlated with age. Using this dataset, we found that ∼108 NPs were expressed by each brain region on average (Table S9); the top brain regions that demonstrated a decrease in NP expression with aging were the hippocampus (25), anterior cingulate gyrus (28), frontal cortex (16), and amygdala (16) (Table S9). We confirmed that the larger decrease of NP expression in these brain regions with aging was not attributed to their intrinsic capacity to express NPs; neither the number nor the average transcript count for NPs differed significantly among surveyed regions (Table S9). A closer examination of the NPs that demonstrated a decrease in expression with age in the aforementioned brain regions revealed that these NPs consisted of expressed ADNPs included in our previous study.^8^ To support the specificity of the observed change in ADNP expression, we analyzed the counts of all non-ADNP NPs in the brain regions under examination and found no age-related changes in their overall levels (Table S10). In short, the spatiotemporal transcriptome analysis presented here showed that the decrease of NPs in the aging human brain was specific to brain regions and NPs implicated in AD, supporting that age-related cognitive decline shares mechanisms with AD and may be mediated by loss of ADNP expression during aging.

**Figure 3.**
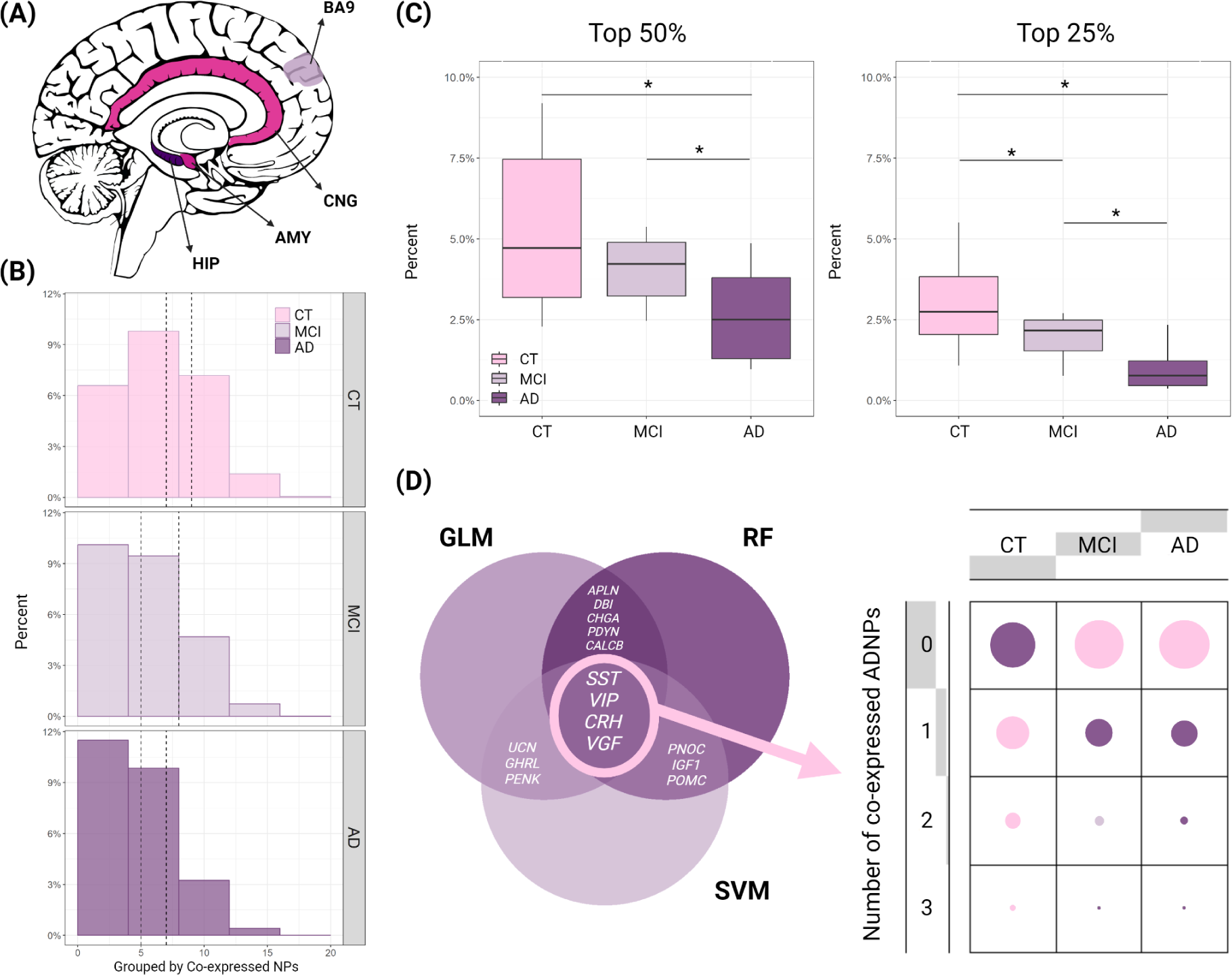
Decreased expression of Alzheimer’s disease-associated neuropeptides (ADNPs) across brain regions during aging as well as cognitive continuum. **(A)** Brain regions (medial sagittal view) showing decreased Alzheimer’s disease-associated neuropeptides (ADNPs) with aging. AMY, amygdala; CNG, anterior cingulate cortex; HIP, hippocampus; BA9, Brodmann area 9. **(B)** Distribution of neurons based on the number of co-expressed ADNPs from the entorhinal cortex (EC) of control (CT), mild cognitively impaired (MCI), and AD donor brains. Median and third quantile of the number of co-expressed ADNPs were marked by dashed lines for each condition. **(C)** The number of neurons above the median and the third quantiles were plotted for CT, MCI, and AD. The decreasing trend of ADNP-co-expressing neuronal count in the cognitive continuum was tested by Jonckheere-Terpstra test. The post-hoc comparisons were tested by Yuen’s two-sample t-test. **(D)** Combination of the neuropeptide outputs (key ADNPs: SST, VGF, VIP, and CRH) derived from three previously reported machine learning algorithms (color modification) ^8^ was used to examine the the number of expressed key ADNPs, ranging from 0 to 3, in the EC of CT, MCI, and AD donor brains. GLM, General Linear Model; SVM, Support Vector Machine; RF, Random Forest; *SST*, Somatostatin gene; *VGF*, Nerve Growth Factor Inducible gene; *VIP*, Vasoactive Intestinal Peptide gene; CRH, Corticotropin-Releasing Hormone gene.

**Table 1.** Significantly decreased expression of ADNPs with aging only occurred in early AD-impacted regions.

| Brain Region | Sample Size | Correlation | p-value | Significance |
| --- | --- | --- | --- | --- |
| Hippocampus | 103 | -0.30 | 0.00 | * |
| Frontal cortex (BA9) | 116 | -0.35 | 0.00 | * |
| Anterior cingulate cortex (BA24) | 88 | -0.30 | 0.01 | * |
| Amygdala | 73 | -0.30 | 0.02 | * |
| Hypothalamus | 109 | 0.02 | 0.85 |  |
| Caudate basal ganglia | 146 | -0.08 | 0.28 |  |
| Nucleus accumbens basal ganglia | 143 | 0.00 | 0.96 |  |
| Putamen basal ganglia | 126 | 0.11 | 0.25 |  |
| Substantia nigra | 66 | -0.23 | 0.08 |  |
| Cerebellar hemisphere | 141 | -0.03 | 0.74 |  |
| Cerebellum | 156 | -0.08 | 0.34 |  |

ADNP, Alzheimer’s Diseases (AD)-associated neuropeptides; BA, Brodmann area; \*, p\_value<0.05, $\alpha = 0.05$ .

To further investigate whether loss of HNP neurons co-expressing ADNPs could be associated with cognitive impairment in early AD, we postulated that brains from individuals with MCI would maintain more of these neurons than AD but not CT donors. Using single-cell transcriptome data generated from CT, AD, and MCI EC region (see Methods), we first confirmed that the correlation between the transcript level of NPs and the number of co-expressed NPs still exist in this dataset (rho = 0.89, p < 0.0001; Figure S2). We subsequently visualized and analyzed the distribution of neurons across three conditions, based on the number of co-expressed neuropeptides (NPs), and marked the median and the 3rd quantile (Q3) values for each condition (Table S11; Figure 3B). Consistent with previous reports (Figure 1B),^8^ distribution of AD neurons showed right skewness in comparison to CT (Figure 3B). As expected, MCI neurons also showed right skewness but less so than that in AD (Figure 3B). To quantify the observation, we analyzed the abundance of neurons above the median ADNP co-expression levels observed in CT for all three conditions; there was a statistically significant decreasing trend in the cognitive continuum from CT, MCI, to AD (p < 0.05; Figure 3C) and CT ECs had significantly higher neuronal counts than MCI and AD (Figure 3C). We furthered the comparison to the third quantile of CT; similarly, the decreasing trend persisted (p < 0.01; Figure 3C) and there were significant differences across these conditions.

Considering subsets of HNP neurons as a representative example, we utilized a combination of the neuropeptide outputs (key ADNPs: *SST, VGF, VIP*, and *CRH*) derived from three previously reported machine learning algorithms (Figure 3D; see Methods).^8^ For neurons in each condition among CT, MCI, and AD, we recorded the number of expressed key ADNPs, ranging from 0 to 3. For neurons expressing no key NPs, no significant differences were observed between MCI and AD but both were more abundant than CT (p < 0.0001; Figure 3D; Table S12&13). For the number of neurons expressing one or three key NPs, significant differences were again only observed where neuronal counts were higher in CT than the other two conditions (p < 0.0001; Figure 3D; Table S12&13). For neurons expressing two key NPs, neuronal counts were highest in CT, followed by MCI, and then AD, with significant differences observed between the groups (p < 0.0001; Figure 3D; Table S12&13). Overall, our analysis supported the early involvement of ADNP expression in AD pathogenesis.

### Physiological distribution of AHNP neurons may mediate brain region vulnerability to AD

“Epicenters” have been described as brain regions showing the most significant pathological alterations, hypothesized to be the initial site of disease onset.^3,4^ Stressed “nodes” are known as brain regions with high network traffic, also referred to as “hubs,” that experience activity-induced deterioration that can lead to or exacerbate diseases.^5,38^ While both concepts are instrumental in theories of AD etiology devised by connectome and network-based studies,^3–5,38^ the cell types underlying these “epicenters” and stressed “nodes” are unclear. Based on the regions that display reduced ADNP expression with aging, we propose that AHNP neurons serve as one of the cellular components of the “hubs” and “epicenters” leading to the onset and/or progression of AD. To test our hypothesis, we analyzed the distribution of AHNP neurons across various regions of the human brain. Two potential observations and implications exist: 1) AHNP neurons ubiquitously exist in all brain regions but those in early-AD impacted regions are, regardless of the underlying cause, more susceptible to dysfunction than others, or 2) AHNP neurons are preferentially distributed in early-AD impacted brain regions to physiologically perform cognitive functions, but they are more prone to dysfunction than other cell types, therefore mediating the regional vulnerability for AD with aging-related cell dysfunctions. We predict that a single-cell transcriptomic survey of brain regions would show that AHNP neurons are more abundantly distributed in early AD-affected brain regions that engage extensively in memory and executive functions, such as the entorhinal cortex, hippocampus, and basal forebrain^39–41^. However, if the first scenario were true, our hypothesis could be negated altogether.

A recent study published single-cell transcriptome data from ∼100 dissections across the forebrain, midbrain, and hindbrain of human donors, and classified brain cells into 461 clusters.^22^ The authors also compared their cell clusters with previous publications that used NP diversity to classify neurons.^22,23^ We first investigated the top brain regions where AHNP neurons using cluster annotations provided by Siletti et al and calculated the number of neurons in each top three brain regions and microdissections (see Methods).^22^ As anticipated, the amygdala and hippocampus—where the age-associated decrease of ADNP expressions was observed^8^—were among the top five brain regions (Figure 4A; Table 2). While existing literature identified the hypothalamus as considerably relevant in AD,^42,43^ it didn’t show an age-related decrease in ADNPs. We also found that the cerebral cortex contained the most AHNP neurons, but this was likely due to the number of dissections assigned to the cerebral cortex. To overcome this issue, we set out to identify the distribution of AHNP neurons among cortical regions separately and ranked the cortical dissections based on the number of AHNP neurons. Again, we found that medial and lateral EC (MEC and LEC) ranked among the top five cortical regions (Figure 4A; Table 3). In particular, the MEC harbored the highest number of AHNP neurons among all micro-dissected regions (Table S14). In contrast, very few AHNP cells were found in the cerebellum, spinal cord, and medulla, where it is generally thought to be spared by AD neuropathology (Table S15), further supporting our hypothesis that dysfunction of AHNP cells mediates the brain region-specific vulnerability to AD.

**Figure 4.**
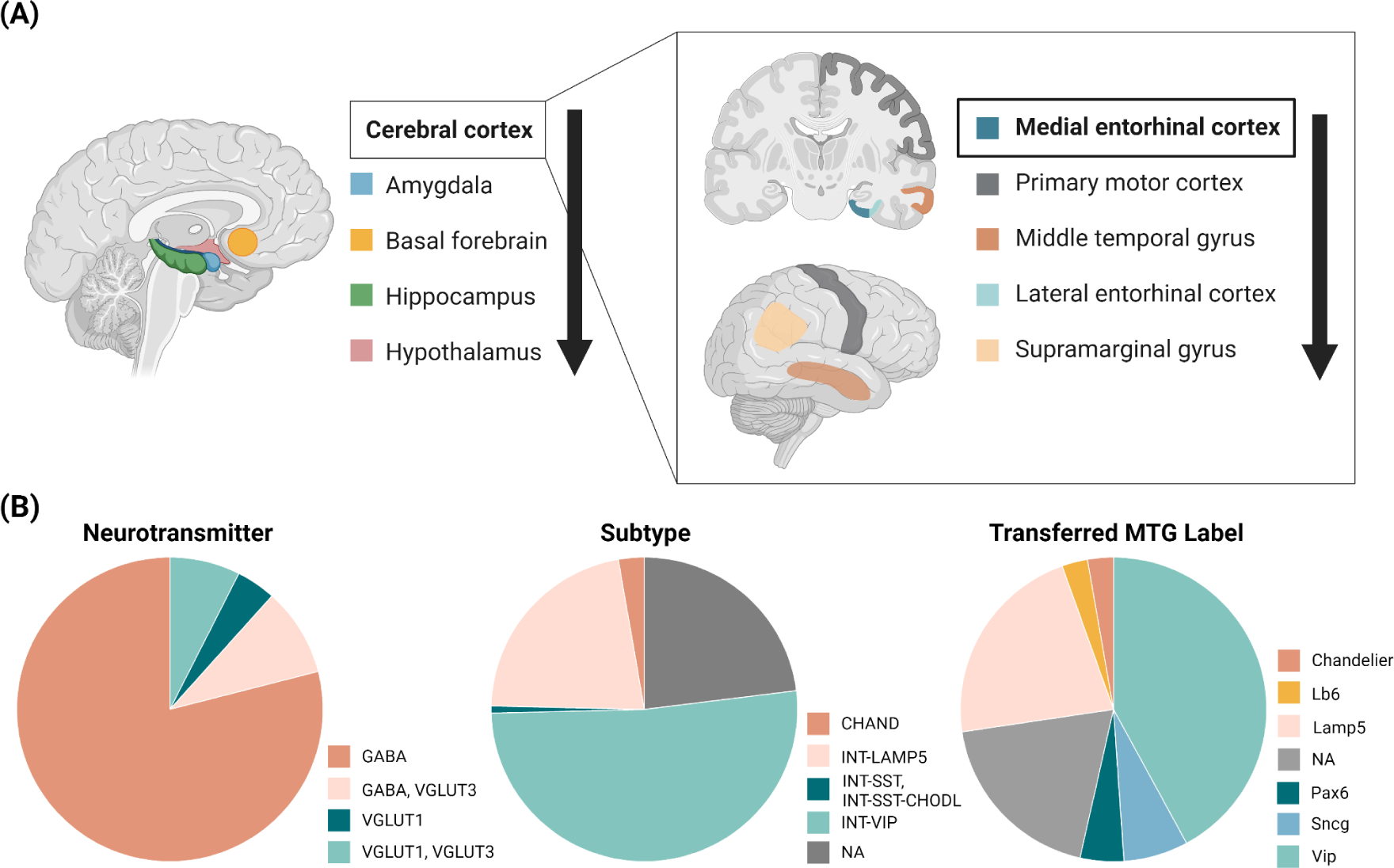
Local density of ADNP-co-expressing high NP-producing (AHNP) neurons may govern brain vulnerability to AD. **(A)** Top five brain and cortical regions ranked by AHNP neuron abundance are visualized. **(B)** Categorization of AHNP neurons predominantly found in EC by neurotransmitter, subtypes, and transferred MTG labels (common cell type nomenclatures for the medial temporal gyrus of the mammalian brain).^22^

**Table 2.** Top five brain regions ranked by AHNP neuron abundance.

| Brain regions | AHNP neuronal count |
| --- | --- |
| Cerebral cortex | 218145 |
| Amygdala | 72256 |
| Basal forebrain | 45559 |
| Hippocampus | 40870 |
| Hypothalamus | 23217 |

**Table 3.**
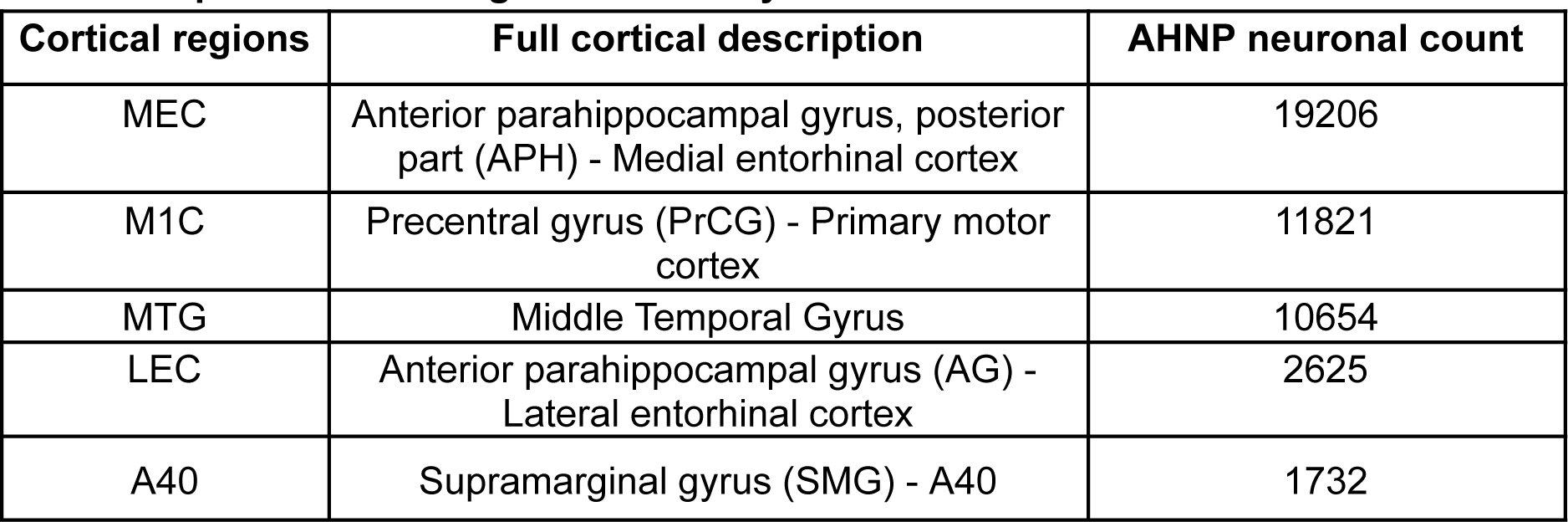
Top five cortical regions ranked by AHNP neuron abundance.

| Cortical regions | Full cortical description | AHNP neuronal count |
| --- | --- | --- |
| MEC | Anterior parahippocampal gyrus, posterior part (APH) - Medial entorhinal cortex | 19206 |
| M1C | Precentral gyrus (PrCG) - Primary motor cortex | 11821 |
| MTG | Middle Temporal Gyrus | 10654 |
| LEC | Anterior parahippocampal gyrus (AG) - Lateral entorhinal cortex | 2625 |
| A40 | Supramarginal gyrus (SMG) - A40 | 1732 |

As the EC—the earliest and most heavily affected region in AD neuropathology—harbored the most significant population of AHNP neurons, we sought to investigate the specific cell types as previous investigation has suggested that most ADNP co-expressing neurons were GABA-ergic.^8^ Specifically, we aimed to understand their how they align with established neuronal cell types. We selected neurons that had MEC in the top three dissections and examined the proportions of neuron types, including based on the neurotransmitter, subtypes, and transferred MTG labels (common cell type nomenclatures for the MTG regions of the mammalian brain).^22^ We found that most of the AHNP neurons were indeed GABA-ergic interneurons; however, about a quarter of the neurons have not been described before (Figure 4B).

## Discussion

Why some ubiquitously expressed proteins (e.g., tau and Aβ) exhibit selective accumulation in particular regions of the brain and cells, yet sparing their comparable neighbors, is fundamental to AD research. Examining multiscale and spatiotemporal RNA-seq data from 1890 human brain samples, we aimed to gain a comprehensive understanding of the potential mechanistic roles that high NP-producing (HNP) neurons that co-express AD-associated neuropeptides (ADNPs) play in mediating the selective vulnerability of brains to AD.

To address our previous hypothesis that HNP neurons, given their secretory and peptidal signaling functions, would demand more translation and transportation, leading to increased metabolic vulnerability,^8^ we investigated the specific characteristics of HNP neurons to the link between HNP neuron dysfunction and the hallmark molecular indicators of AD. We found that 1) HNP cells were more metabolic active and had gene expression profiles suggesting higher connectivity; 2) MEC AD neurons co-expressing higher levels of NPs showed the molecular signature of AD, including protein misfolding; and 3) the deficiency of AD cells was linked to loss-of-functions of HNP cells. While we anticipated a greater metabolic burden as a source of neuronal vulnerability for HNP neurons, it was surprising to discover that HNP neurons can be predisposed to tau hyperphosphorylation and misfolding.^27,44^ In addition to *ERBB4*, recent research has revealed that the impairment of NMD, an elevated process observed in HNP cells and suppressed by cellular stress, mediates tau-induced neurotoxicity.^45^ Therefore, we posit that the disruption of cellular processes elevated in HNP neurons could occur more readily/earlier in these cells and contribute to the formation of misfolded proteins within them, subsequently leading to their selective degeneration. This position aligns with the existing knowledge that brain regions with strong anatomical connectivity harbored more misfolded proteins^37,38^; the cell clusters carrying out the functions, such as HNP neurons with greater connectivity and metabolic activity described in this study, are likely more prone to protein misfolding–the general framework of which was already proposed first as “nodal stress” and explained in detail by others.^5,6,38^ Further, it is generally agreed that brain regions accumulating most pathology-associated proteins (disease-susceptible brain regions) physiologically contain the cells that are most susceptible to cytotoxic events and are among the earliest affected in the disease’s progression.^3,4,6^ Taken together, at the cellular level, our results indicate that disproportionate loss of HNP neurons in AD could be a reflection of their earlier involvement in the disease process as the result of higher chances of cytotoxic events–which are associated with their functions–occurring, such as propensity to oxidative stress, mitochondrial dysfunction, and accumulation of misfolded proteins.

To comprehensively demonstrate that the loss of ADNPs expressed by HNP neurons participates in early AD development and/or progression, we considered perspectives from both aging and MCI. We analyzed spatiotemporal transcriptomic data of brains from the natural aging populations generated by the GTEx consortium to reveal that the decrease in ADNP expressions with aging is specific to early AD-associated brain regions. Supporting our hypothesis that dysfunction of ADNP expression in HNP (AHNP) neurons appears early in AD, we observed a significant decline in the abundance of AHNP neurons across the continuum from CT, progressing through MCI, to AD. These results are consistent with previous studies that underscored the role of key ADNPs play in processes crucial to the pathogenesis of neurodegeneration, such as mitochondrial dysfunction, persistent neuroinflammation, and disrupted circadian rhythm (briefly summarized by Li and Larsen),^8^ indicating that alterations in the abundance and diversity of ADNP-producing neurons likely participate in the onset and/or progression of cognitive decline and neurodegeneration in AD. Our findings also highlight the relevance of ADNPs as combinatorial and longitudinal biomarkers to evaluate the risk and progression of AD development. For instance, an analysis of the CSF proteome identified that CHGA and VGF exhibited significant differences in abundance among CT, MCI, and AD groups.

Further studies validating these findings and investigating the underlying mechanisms responsible for the observed decline in AHNP neuron abundance are needed. Evaluating whether interventions aimed at mimicking and/or preserving the ADNP signaling network could impede progression of AD is also valuable, especially considering the subtle differences in GPCR expression observed between CT and AD.^8^ Collectively, these observations suggest that deficiencies of ADNPs with aging contribute to AD development and progression and that these deficits could be a consequence of losing AHNP neurons during aging, which underlies shared cognitive decline during aging and in early AD.

Cellular dysfunctions, characterized by distinct molecular pathways and biochemical properties, play crucial roles in the onset and progression of AD.^6^ Extensive research over the past decade has delved into connectome-based studies of neurodegenerative diseasesand and provided profound understanding of the brain regions and neural circuitries vulnerable to AD^3,4,7,38^; however, the specific cell types and cellular components that underpin these connectomic findings in relation to AD remain elusive. Therefore, our presentation of the physiological distribution of HNP neurons within the brain is fundamental to deciphering their roles in AD; their preferential localization in AD-susceptible brain regions further signifies their potential role as cellular elements contributing to the initiation of AD. However, potential confounding variables, such as transcription factors and epigenetic mechanisms, warrant careful consideration. Recent research has made tremendous advancements on providing a comprehensive multi-omic brain atlas for physiological conditions,^22,46,47^ our results suggest that such efforts will also be vital to explore brains of diseased states. By understanding how dysfunction in HNP neurons occurs with aging and its detrimental effects on interconnected behavioral domains, we can gain valuable insights into the cell types and dysfunctions at the early stages of AD to develop effective disease models as well as cell type-specific targeting strategies for prevention and therapy.

## Consent Statement

Consent from human subjects was not necessary.

## Funding

ML is supported by PAL’s discretionary funds and the Doctoral Dissertation Fellowship from the Graduate School Fellowship Office at the University of Minnesota.

## Author contributions

ML conceived the study. ML developed and performed bioinformatic analyses and data visualization. ML and PAL discussed the results. ML wrote the manuscript. ML and PAL revised the manuscript. NF validated the bioinformatic analysis and verified reproducibilty of results and outputs. All authors approved the final manuscript.

## Acknowledgments

Dr. Alice Larson provided helpful comments and suggestions that strengthened this manuscript. The data used for the analyses described in this manuscript were obtained from the GTEx Portal in October 2022. The Genotype-Tissue Expression (GTEx) Project was supported by the Common Fund of the Office of the Director of the National Institutes of Health and by NCI, NHGRI, NHLBI, NIDA, NIMH, and NINDS. The results published here are in whole or in part based on data obtained from the AD Knowledge Portal. The data available in the AD Knowledge Portal would not be possible without the participation of research volunteers and the contribution of data by collaborating researchers. The results published here are in whole or in part based on data obtained from the AD Knowledge Portal (https://adknowledgeportal.org/). Study data were generated from postmortem brain tissue provided by the Religious Orders Study and Rush Memory and Aging Project (ROSMAP) cohort at Rush Alzheimer’s Disease Center, Rush University Medical Center, Chicago. This work was supported in part by the Cure Alzheimer’s Fund, NIH grants AG058002, AG062377, NS110453, NS115064, AG062335, AG074003, NS127187, MH119509, HG008155 (M.K.), RF1AG062377, RF1 AG054321, RO1 AG054012 (L.-H.T.) and the NIH training grant GM087237 (to C.A.B.). ROSMAP is supported by P30AG10161, P30AG72975, R01AG15819, R01AG17917. U01AG46152, U01AG61356. ML is supported by the Doctoral Dissertation Fellowship granted through the Graduate School Fellowship Office at the University of Minnesota. PAL also provided discretionary funds supporting ML. Figures were organized using BioRender (biorender.com).

## Conflict of interest statement

All authors declare no conflict of interest.

## Supporting information

Supplementary materials for this manuscript can be found at https://github.com/mancili/HNP.

## Notes

https://github.com/mancili/HNP

### Competing Interest Statement

The authors have declared no competing interest.

### Summary of Updates

Add coauthor, Dr. Nicole Flack, who contributed significantly to validate the reproducibility and quality of the manuscript

